# Eco-evolutionary dynamics create extinction avalanches in mutualistic-antagonistic networks

**DOI:** 10.64898/2026.09.21.753184

**Authors:** Felix Jäger, Christian Guill, Nicolas Loeuille, Youssef Yacine, Korinna T. Allhoff

## Abstract

Evolution has given rise to the diversity of life on Earth, but it can also cause species extinctions. While major extinction events are usually associated with external drivers, in this study, we show mechanistically how evolution can trigger abrupt extinction avalanches in ecological networks without any environmental change. In particular, we investigate an eco-evolutionary simulation model of a tripartite mutualistic-antagonistic network, such as a plant-pollinator-herbivore network, with additional intraguild competition. The prevalence of self-organised extinction avalanches depends on a subtle balance of the different interaction types, where antagonism and similarity-based competition are needed for initial diversification while mutualism and trait-independent competition foster evolutionarily driven extinction events. Extinction avalanches are triggered by a mutualist evolving to interact with plants that are not subject to sufficient antagonistic control. Subsequently, the mutualist, its plant partners and their main antagonist potentially outcompete the rest of the community. Our findings point out that evolutionary murder may claim many victims simultaneously and highlight the need to take different interaction types into account to gain a comprehensive understanding of the links between evolution and biodiversity in ecological communities.

## Introduction

Major extinction events are an integral part of the evolution of ecological communities across scales. To date, five big macroevolutionary mass extinctions have shaped Earth history, and explaining their causes has been a central objective in macroevolution and paleobiology (Hallam and Wignall, 1997; Newell, 1967; Raup, 1986). However, extinction avalanches are also observed on smaller spatial and temporal scales, such as in local or regional ecological networks, with inevitable consequences for biodiversity and ecosystem functioning (Brook et al., 2003; Donohue et al., 2017; Estes et al., 2016; Paine, 1966; Sanders et al., 2013). Understanding why and under which conditions such extinction events occur is therefore of utmost importance for evolutionary biology, community ecology and conservation biology.

Usually, major extinction events, i.e., the loss of a large proportion of the diversity within a system over a short period of time, are associated with external drivers that act as potential triggers. For instance, continuous changes in atmospheric CO_2_ levels, or specific events such as large scale volcanic activity are hypothesised drivers of mass extinctions (Bond and Grasby, 2017). At local to regional scales, biological invasions can induce extinctions (Bellard et al., 2016; Burbidge and Manly, 2002; Miller et al., 1989), and gradually increasing insect mortality may cause the abrupt collapse of pollination networks (Dakos and Bascompte, 2014; Lever et al., 2014).

Another interesting possibility is for extinction avalanches to be self-organised (Bak and Sneppen, 1993). By this we mean that the extinctions emerge only from the intrinsic eco-evolutionary dynamics of the community without any external forcing. Most often, evolution is associated with creating biodiversity, e.g., via adaptive radiations following key innovations (Futuyma and Agrawal, 2009; Miller et al., 2023; Weber and Agrawal, 2014), or, at microevolutionary scale, through the creation and maintenance of polymorphisms (Apanius et al., 1997; Rozen and Lenski, 2000). Some empirical studies, however, have shown that evolution can also drive extinctions (Fiegna and Velicer, 2003; Greenrod et al., 2026; Olsen et al., 2004), and it is hypothesised that evolution may accelerate or even trigger ecosystem tipping points (Ardichvili et al., 2023; Dakos et al., 2019). In the theoretical literature, the process where evolution of one species can entail extinctions of others has been termed ‘evolutionary murder’ (Leoz et al., 2026; Parvinen, 2005; Shang et al., 2024; Weinbach et al., 2022; Weyerer et al., 2023), and it has been recognised that evolutionary murder can play an important role in the context of ecological networks (Loeuille, 2019).

An open question is how different types of interactions, such as mutualism, antagonism or competition, contribute to evolution-driven extinction events (Leoz et al., 2026; Loeuille, 2019). Recent studies in the context of evolutionary murder focus on single interaction types and suggest that competition and mutualism promote evolutionary murder (Leoz et al., 2026; Loeuille, 2019; Weinbach et al., 2022; Weyerer et al., 2023), while results for antagonism are ambiguous (Shang et al., 2024; van Velzen, 2023). Real-world ecological communities, however, usually include multiple types of interaction simultaneously (Hackett et al., 2019; Kéfi et al., 2015; Melián et al., 2009; Morrison et al., 2020; Pocock et al., 2012). The additional indirect effects arising in such networks are hypothesised to have intricate implications for the eco-evolutionary dynamics (Fontaine et al., 2011), and could open up new pathways for evolution to cause extinctions.

Here, we aim to unravel the mechanisms behind recurrent extinction avalanches in communities that combine different types of interaction. We use an eco-evolutionary simulation model of tripartite mutualistic-antagonistic networks such as plant-pollinator-herbivore networks that also include intraguild competition. In a previous article, Jäger et al. (2026c) used a version of the model to examine diversification patterns and also reported the occurrence of extinction avalanches, but did not investigate them systematically. These avalanches represent an intriguing example of self-organised extinction events that arise solely from a mutation-selection process and the involved biotic interactions. They can hence be interpreted as an extreme form of evolutionary murder, where evolution in some species makes not just one victim, but many. In this study, we ask (1) which conditions determine size and frequency of extinction avalanches and (2) what is the eco-evolutionary mechanism underlying extinction avalanches in the model. Ultimately, we improve our understanding of how different types of ecological interactions contribute to the influence of evolution on the (in)stability of ecological communities (Loeuille, 2010).

## Methods

We model the evolution of a tripartite ecological network, consisting of plants, mutualists and antagonists, based on a previous model version by Jäger et al. (2026c). Starting with only one phenotype in each of the guilds, the model allows for subsequent diversification and possibly for recurrent extinction avalanches. In this section, we first describe the community dynamics in the model, followed by the evolutionary dynamics. Finally, we explain the analyses performed on the simulation output. We refer to Jäger et al. (2026c) for more detailed explanations on the underlying model assumptions. All code and data are available online (Jäger et al., 2026a).

### Community dynamics

The model includes three types of biotic interactions: (i) mutualism between the mutualist and the plant guild, (ii) antagonism between the plant and the antagonist guild and (iii) additional intraguild competition within each of the guilds. We refer to the members of the guilds as ‘phenotypes’. The density dynamics of the phenotypes in the system are described by the following differential equations:

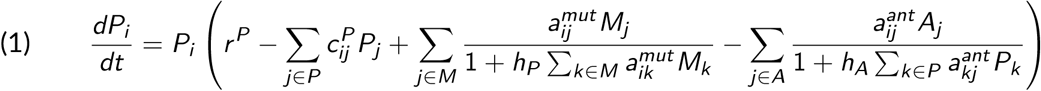

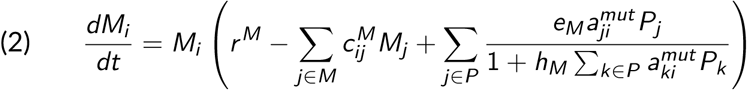

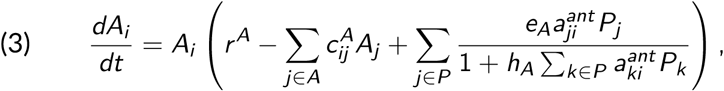

where *P_i_* (*M_i_*, *A_i_*) denotes the density of a plant (mutualist, antagonist) phenotype *i*. *r ^P^* (*r ^M^*, *r ^A^*) is the net intrinsic growth rate of plants (mutualists, antagonists) in the absence of biotic interactions. Note that we assume *r ^P^ >* 0, reflecting that plants are autotrophs; however, *r ^M^* and *r ^A^* are negative, hence the interaction with plants is obligate for the mutualists and antagonists. Plant (mutualist, antagonist) phenotypes *i* and *j* compete with each other with competition strength 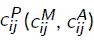. Interguild mutualism and antagonism terms follow a Holling type II functional response with saturation coefficients *h_P_*, *h_M_* and *h_A_*for plants, mutualists and antagonists, respectively, corresponding to handling times in classical predator-prey models. The strength of mutualism (antagonism) between a plant phenotype *i* and a mutualist (antagonist) phenotype *j* is denoted by 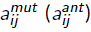. *e_M_* and *e_A_*denote conversion efficiencies of mutualists and antagonists, respectively.

The strengths of the mutualistic and antagonistic interactions in the model depend on traits of the involved phenotypes. Each phenotype *i* possesses a single quantitative trait *q_i_*. The strength of interguild mutualism (antagonism) between a plant phenotype *i* and a mutualist (antagonist) phenotype *j* increases with the similarity of their traits, as captured by Gaussian functions:

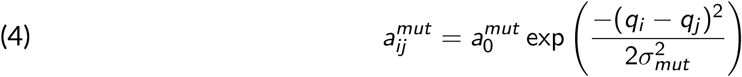

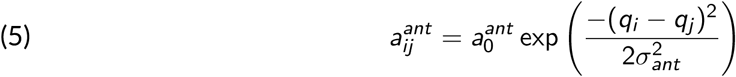

The parameter 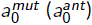 is the maximum mutualistic (antagonistic) interaction strength, while the interaction kernel widths *σ_mut_*and *σ_ant_*capture how fast interactions become weak when partner traits differ. Note that mutualistic and antagonistic interactions are mediated by the same plant trait, which represents a case of ecological pleiotropy and favours simultaneous diversification of all guilds in the model (Jäger et al., 2026c).

Intraguild competition between phenotypes *i* and *j* consists of two components. Some of the competition is based on trait matching, similar to the interguild interactions. This can, for instance, reflect direct interference competition. The rest of the competition is trait-independent, e.g., reflecting competition for external resources, space or nesting sites. Total competition is a combination of these two components:

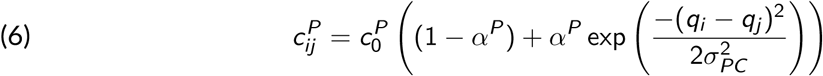

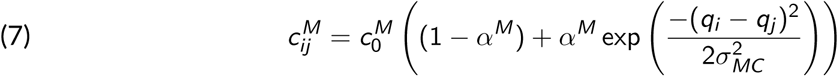

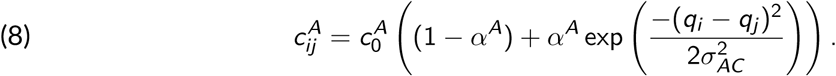

Here, 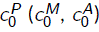 is the maximum competition strength for plants (mutualists, antagonists); the widths of the competition kernels *σ_PC_*, *σ_MC_* and *σ_AC_* describe how fast competition gets weaker for dissimilar phenotypes. *α^P^*, *α^M^* and *α^A^* denote the proportion of competition that is similarity-based as opposed to trait-independent.

### Evolutionary dynamics

Evolution is modelled via a mutation-selection process. Simulations start with only one phenotype in each guild with small initial density *N*_0_; all traits are initialised to 0. The algorithm then proceeds by simulating community dynamics and mutation events in turn. For the community dynamics, the differential equations 1-3 are solved numerically. At a mutation event, a parent phenotype is selected at random among all guilds, with probability proportional to the product of its density and the respective mutation rate of an individual (*µ_P_*, *µ_M_* or *µ_A_*for plants, mutualists, antagonists). Then, an offspring phenotype is created whose trait value is drawn from a normal distribution around the parent trait with small standard deviation Σ. Its initial density is *N*_0_, subtracted from the parent’s density. At each mutation event, all phenotypes with density below *N*_0_ are considered extinct and removed from the system.

At each mutation event, the waiting time until the next mutation is drawn from an exponential distribution with parameter *µ_P_P_tot_* + *µ_M_M_tot_* + *µ_A_A_tot_*, where *P_tot_*, *M_tot_* and *A_tot_* denote total densities of plants, mutualists and antagonists. This reflects that mutations follow a Poisson process. The time of the first mutation event is exempt from this rule due to the low initial densities and is set to 1 000 time units.

### Model analysis

Simulations were run for 5 billion time units or until the number of phenotypes in at least one of the guilds exceeded the threshold of 300, to avoid excessive runtimes (200 when varying *α^P^*, *α^M^*and *α^A^*). We tracked the densities and the trait values of all phenotypes through time. For the analysis of the simulation output, the phenotypes in each guild were grouped into branches, collections of phenotypes with close trait values that can be considered functionally equivalent. In theory, in the long run, all but one phenotype in a branch would go extinct due to competitive exclusion. However, in the simulations (and in nature) new mutations often accumulate faster than it takes for slightly weaker competitors to go extinct. Branches were identified via a clustering analysis using the DBSCAN algorithm, which is a widely used density-based non-parametric clustering algorithm (Ester et al., 1996).

Extinction avalanches were defined as the loss of more than 50% of the branches in a guild, but at least 4 branches, within at most 100 million time units. As branches represent groups of similar phenotypes, this definition focuses on the loss of functional diversity rather than mere species richness, and thus addresses a dimension of biodiversity that is more meaningful in terms of the functioning and stability of ecosystems (McCann, 2000). Our results were qualitatively robust to quantitative changes in the proportion of branches that need to be lost as well as to changes in the maximum time window (see Supplementary Material S1; Jäger et al., 2026b).

To pinpoint the conditions under which extinction avalanches occur, we varied the relative strengths of the different interaction types (including competition modes) and the relative speeds of evolution. Importantly, Jäger et al. (2026c) showed that the system tends to diversify significantly across all guilds only when the relative strength of antagonism and the relative speed of evolution of antagonists is sufficiently high and when some of the competition in the system is similarity-based. Here, we ran simulations with (i) varying maximum strength of mutualism and antagonism (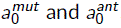, ranging from 0 to 2), (ii) varying relative mutation rates (*µ_M_*: *µ_P_* and *µ_A_*: *µ_P_*, ranging from 0.25 to 4) and (iii) varying proportions of similarity-based competition (*α^P^*, *α^M^* and *α^A^*, ranging from 0 to 1). For each of these sets of simulations, the values of all other parameters were chosen according to the baseline parametrisation introduced in Jäger et al. (2026c) and given in Table 1. For each parameter combination, the rate of extinction avalanches (number of extinction avalanches over total simulated time) and their mean size (number of branches lost) was recorded, using 5 replicates (see Supplementary Material S2 for details on how number and size of extinction avalanches were computed).

**Table 1.** – Model variables and parameters.

| Symbol | Meaning | Default value | Dimension |
| --- | --- | --- | --- |
| **Variables** |  |  |  |
| $P_i, M_i, A_i$ | plant/mutualist/antagonist phenotype densities | - | individuals · area<sup>-1</sup> |
| $q_i$ | plant/mutualist/antagonist trait | - | trait dimension* |
| **Ecological parameters** |  |  |  |
| $r^P$ | plant intrinsic net growth rate | 10 | time<sup>-1</sup> |
| $r^M, r^A$ | mutualist/antagonist intrinsic net growth rate | -0.01 | time<sup>-1</sup> |
| $e_M, e_A$ | mutualist/antagonist conversion efficiency | 3 | none |
| $h_P, h_M, h_A$ | plant/mutualist/antagonist saturation coefficient | 0.1 | time |
| $a_0^{mut}, a_0^{ant}$ | maximum strength of mutualism/antagonism | 1 | area · (individuals · time)<sup>-1</sup> |
| $\sigma_{mut}, \sigma_{ant}$ | mutualism/antagonism kernel width | 2.5 | trait dimension* |
| $c_0^P$ | maximum plant competition coefficient | 10 | area · (individuals · time)<sup>-1</sup> |
| $c_0^M, c_0^A$ | maximum mutualist/antagonist competition coefficient | 1 | area · (individuals · time)<sup>-1</sup> |
| $\alpha^P, \alpha^M, \alpha^A$ | plant/mutualist/antagonist share of similarity-based competition | 0.2 | none |
| $\sigma_{PC}, \sigma_{MC}, \sigma_{AC}$ | plant/mutualist/antagonist competition kernel width | 1 | trait dimension* |
| $N_0$ | extinction threshold/initial mutant density | 10<sup>-5</sup> | individuals · area<sup>-1</sup> |
| **Evolutionary parameters** |  |  |  |
| $\Sigma$ | Mutation amplitude (standard deviation) | 0.05 | trait dimension* |
| $\mu_P, \mu_M, \mu_A$ | Mutation rate of a plant/mutualist/antagonist individual | 2 · 10<sup>-6</sup> | area · (individuals · time)<sup>-1</sup> |

\*The dimension of the traits depends on the study system.

To further understand the mechanism underlying extinction avalanches, we performed manipulative experiments based on observed example avalanches where evolution and/or ecological dynamics of specific branches are turned off. These are explained in detail in the Results section.

## Results

### Extinction avalanches require a balanced interplay of mutualism, antagonism and competition

For extinction avalanches to occur, mutualism and antagonism strength as well as speed of mutualist and antagonist evolution need to be in balance (Fig. 1A and B). If mutualism is much stronger than antagonism or mutualists evolve much faster than antagonists, diversification in the network remains limited in the first place, indicated by dots in Figure 1 (as observed in Jäger et al., 2026c, see also Supplementary Material S3). This constrains the potential for extinction avalanches and explains their low rate in the bottom right parts of Fig. 1A and B. If, on the other hand, mutualism is much weaker than antagonism or mutualists evolve much slower than antagonists, the rate of extinction avalanches is also low (top left regions in Fig. 1A and B), even though diversification is high (cf. Fig. S3). Furthermore, extinction avalanches are more frequent when the overall antagonism and mutualism strength is high (red area in top right part of panel A). Similarly, avalanche rate increases if overall mutualist and antagonist evolutionary speed is high compared to that of plants (top right part of panel B).

**Figure 1.**
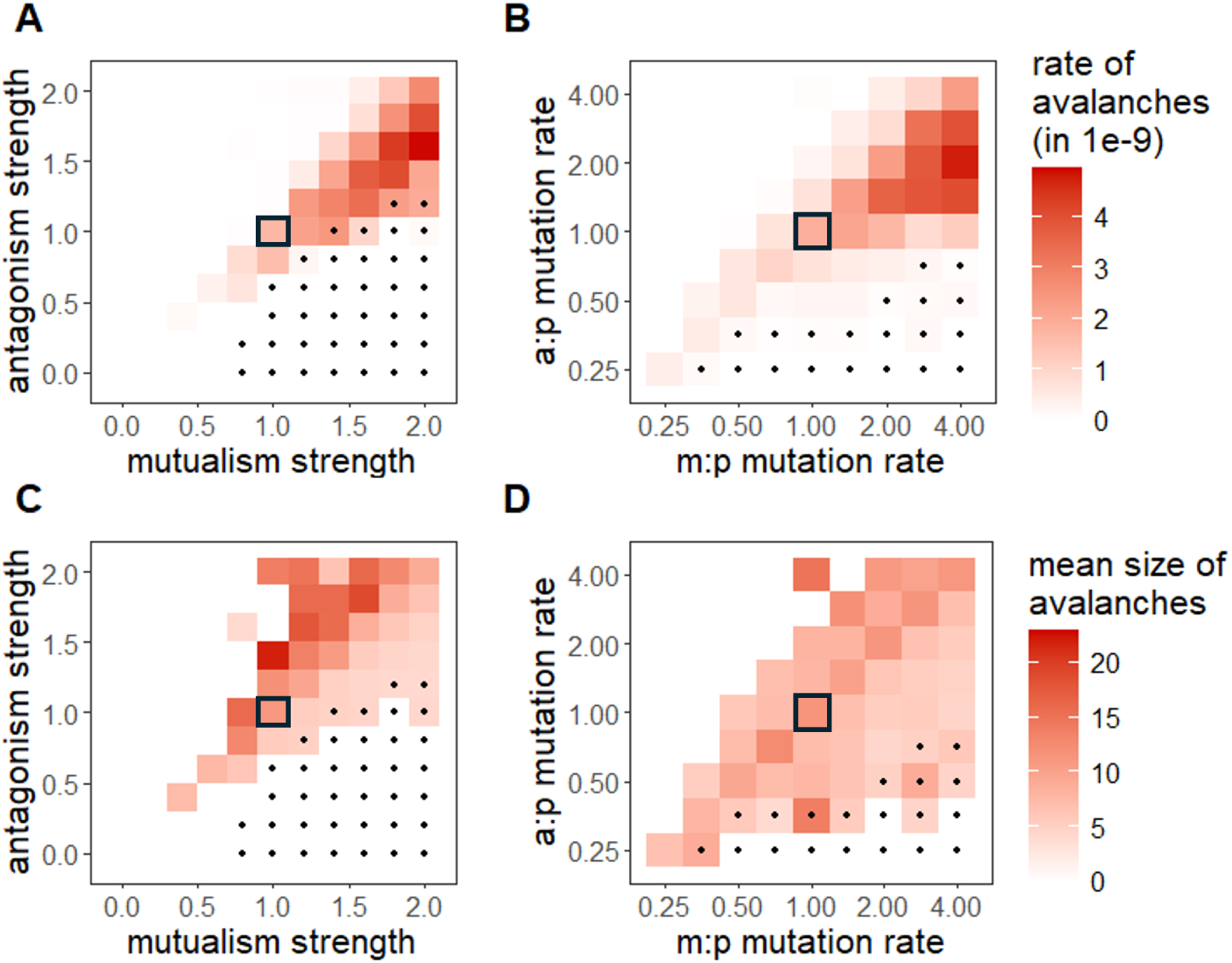
– Prevalence of extinction avalanches under varying antagonism/mutualism strength and varying speed of evolution. (A, B) Rate of extinction avalanches, measured as the total number of extinction avalanches in any of the guilds over total simulated time, using 5 replicates for each parameter combination. Dots indicate low overall diversification, i.e., the maximum total number of branches within a simulation run, averaged across all replicates, is less than 15. (C, D) Size of extinction avalanches, measured as the maximum number of branches lost within at most 100 million time units. A mean is taken over all avalanches in all guilds, using 5 replicates. Rate and size are computed for (A, C) varying maximum strength of mutualism and antagonism (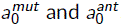), and (B, D) varying relative mutualist and antagonist mutation rates (*µ_M_*: *µ_P_* and *µ_A_*: *µ_P_*; *µ_P_* = 2 *·* 10*^−^*^6^). All other parameters were chosen according to Table 1. The highlighted tiles represent the baseline scenario used for subsequent analyses below.

Within the parameter space where extinction avalanches take place, high avalanche rate is rather associated with small avalanche size (Fig. 1C and D). This can be explained by the fact that if extinction avalanches are set off earlier, the system has not yet diversified much, constraining the size of avalanches. Besides, extinction avalanches in plants tend to be larger than in the other guilds (see Supplementary Material S4). This can be attributed to the fact that plants reach higher levels of diversity in the first place since mutualists and antagonists depend on plants but not vice versa (see Supplementary Material S3).

Extinction avalanches only occur if there is sufficient trait-independent competition, especially in the plants (Figure 2). If *α^P^* = 1, meaning only similarity-based plant competition, avalanches become very rare and possible only for low values of *α^M^* and *α^A^*. For intermediate values of *α^P^*, avalanches tend to occur when *α^M^* is larger than *α^A^*. When there is only trait-independent competition in the plants (*α^P^* = 0) and not enough similarity-based competition in the other guilds, avalanche potential is additionally constrained by limited initial diversification, indicated by dots in the figure (as observed in Jäger et al., 2026c, see also Supplementary Material S3). The results on the sizes of extinction avalanches for varying proportion of similarity-based competition confirm the afore-mentioned correlation of high avalanche rates with small sizes and are reported in Supplementary Material S4.

**Figure 2.**
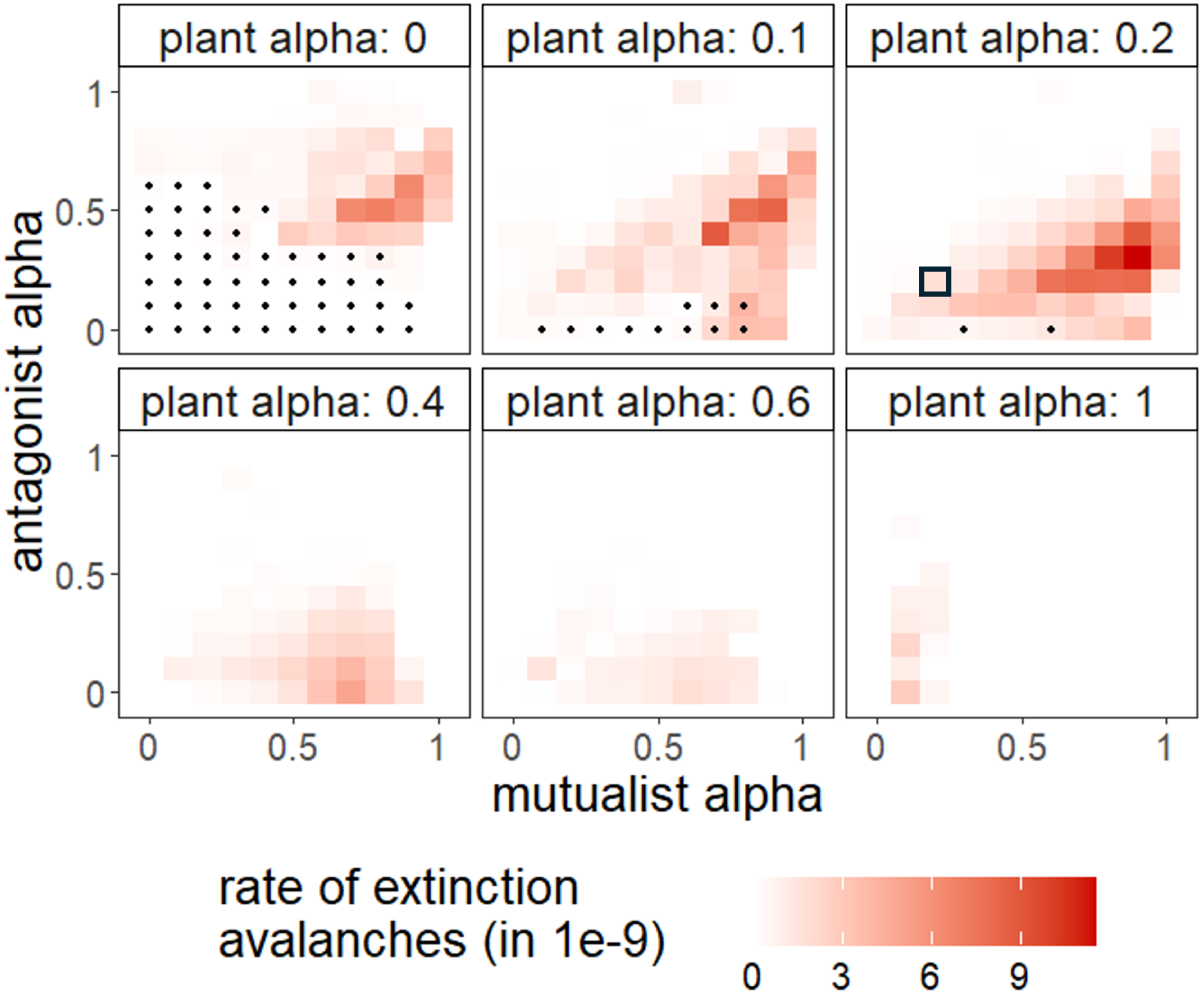
– Prevalence of extinction avalanches under varying proportion of similarity-based competition. Plots show the rate of extinction avalanches measured as the total number of extinction avalanches in any of the guilds over total simulated time. Rate of extinction avalanche is measured using 5 replicates for varying shares of similarity-based competition (*α^P^*, *α^M^*and *α^A^*). All other parameters were chosen according to Table 1. Dots indicate low overall diversification, i.e., the maximum total number of branches within a simulation run, averaged across all replicates, is less than 15. The highlighted tile represents the baseline scenario used for subsequent analyses below.

### The mechanism underlying extinction avalanches

To obtain a mechanistic understanding of the extinction avalanches, which allows for explaining the patterns found in the previous section, we analysed an example avalanche in a simulation run using the baseline parametrisation (Fig. 3). The avalanche exhibits the following sequence of events (see Fig. 3C): First, a large proportion of mutualist branches goes extinct within just a few mutation events, immediately followed by an extinction avalanche in the antagonists. The extinction of the majority of plants occurs later, or not at all (e.g., in the second extinction avalanche in Fig. 3A after around 2.2 billion time units). This sequence of events could be confirmed in 20 replicate runs with the same parametrisation (see Supplementary Material S2).

**Figure 3.**
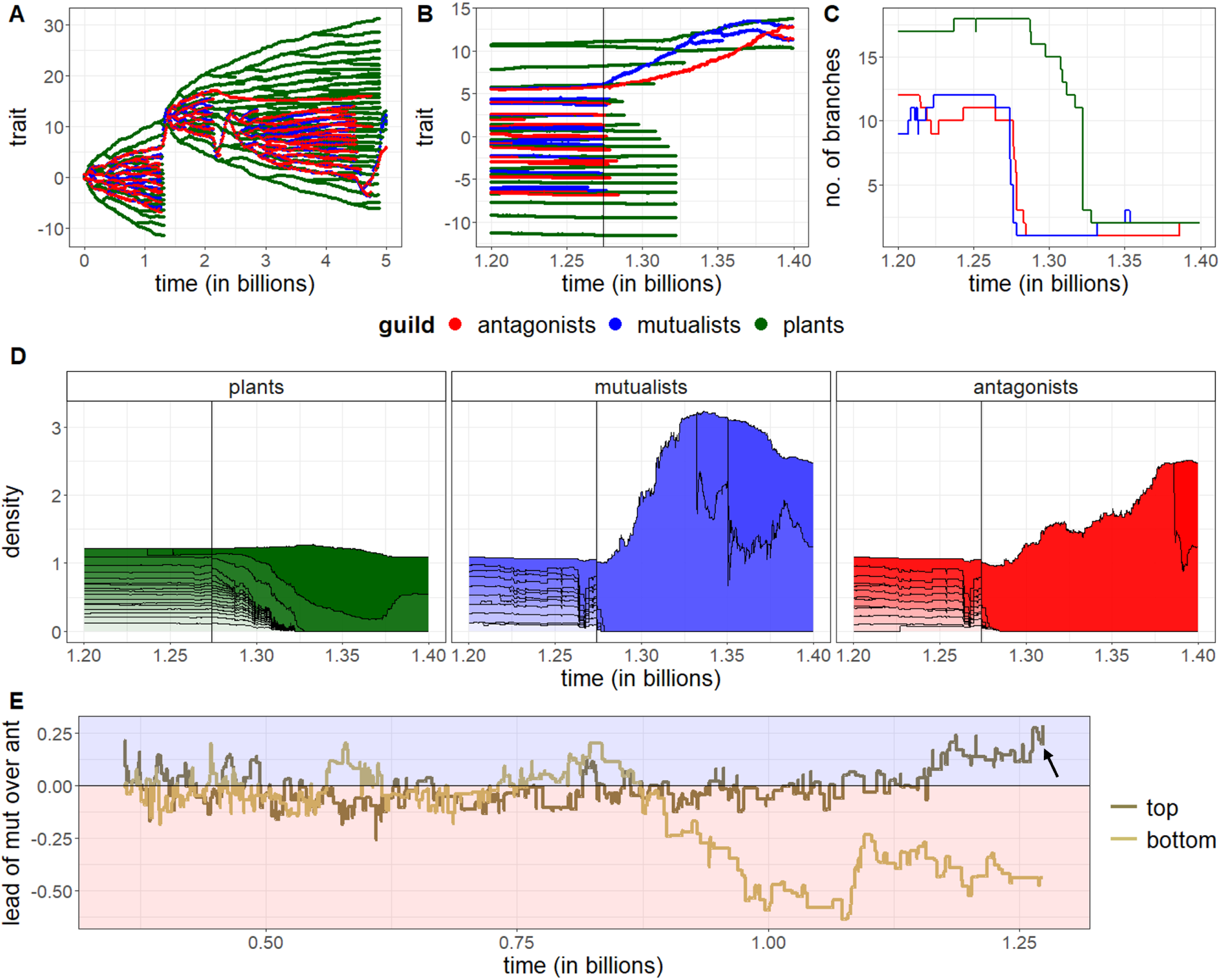
– Example of an extinction avalanche. (A) Full time series showing the traits of all phenotypes. (B) Close-up of the first extinction avalanche in the time series. The vertical black line indicates the point in time right before the beginning of the extinction avalanche, used for subsequent analyses below. (C) Number of branches over time in the focal time window. (D) Densities of branches over time in the focal time window, stacked by guild. Within guilds, branches are ordered by their mean trait value. (E) Difference between the top mutualist and the top antagonist trait value and difference between the bottom antagonist and bottom mutualist trait value over time in the period before the focal extinction avalanche, starting from the last time there were at most 4 mutualist branches. Positive values indicate a more extreme mutualist trait (shaded in blue), negative values a more extreme antagonist trait (shaded in red). All parameters were chosen according to Table 1.

Extinction avalanches are triggered by a mutualist branch approaching outer plant branches ahead of an antagonist. In the period before the example avalanche, the lead of the top mutualist branch over the top antagonist branch in the trait space shows an increasing trend, meaning that the top mutualist gets closer to the outer plant branches, distancing the next antagonist branch (Fig. 3E). This is associated with a growing imbalance in densities within the guilds, in favour of the top mutualist and the top antagonist (Fig. 3D), since the top mutualist gets a competitive advantage by interacting with plant branches without an antagonist close by, while the next antagonist also benefits from increasing densities of its plant resources. Once the lead of the mutualist over the antagonist crosses a threshold, the system ‘tips over’ and the growing imbalance in densities continues into the observed extinction avalanche (black line in Fig. 3D, or end of the plot in Fig. 3E). Note that in the example, shortly before the avalanche, the imbalance in mutualist and antagonist densities is temporarily reduced again (Fig. 3D), which can be attributed to a decrease in the lead of the top mutualist before the final increase (see arrow in Fig. 3E). Such fluctuations in relative densities are likely to precede an extinction avalanche since the densities are particularly sensitive to stochastic changes in the lead when the lead gets close to the tipping threshold. Note that the described process can equally likely happen with the bottom mutualist gaining a lead over the bottom antagonist branch (e.g., in the second extinction avalanche in Fig. 3A after around 2.2 billion time units).

The top mutualist branch in the example avalanche is the evolutionary ‘murderer’, outcompeting the other mutualists (note that there is direct trait-independent competition in all guilds in the model). This is confirmed by manipulative experiments, where right before the extinction avalanche from Fig. 3B the evolution of single branches was turned off (Fig. 4A). Subsequent extinction avalanches are prevented if and only if it is the top mutualist branch (blue triangle in Figure 4A) that does not evolve. Moreover, suppression of the evolution of the top antagonist branch (red triangle) greatly increases the extent of extinction avalanches, as in this case the mutualist easily gains a substantial lead over the non-evolving antagonist.

**Figure 4.**
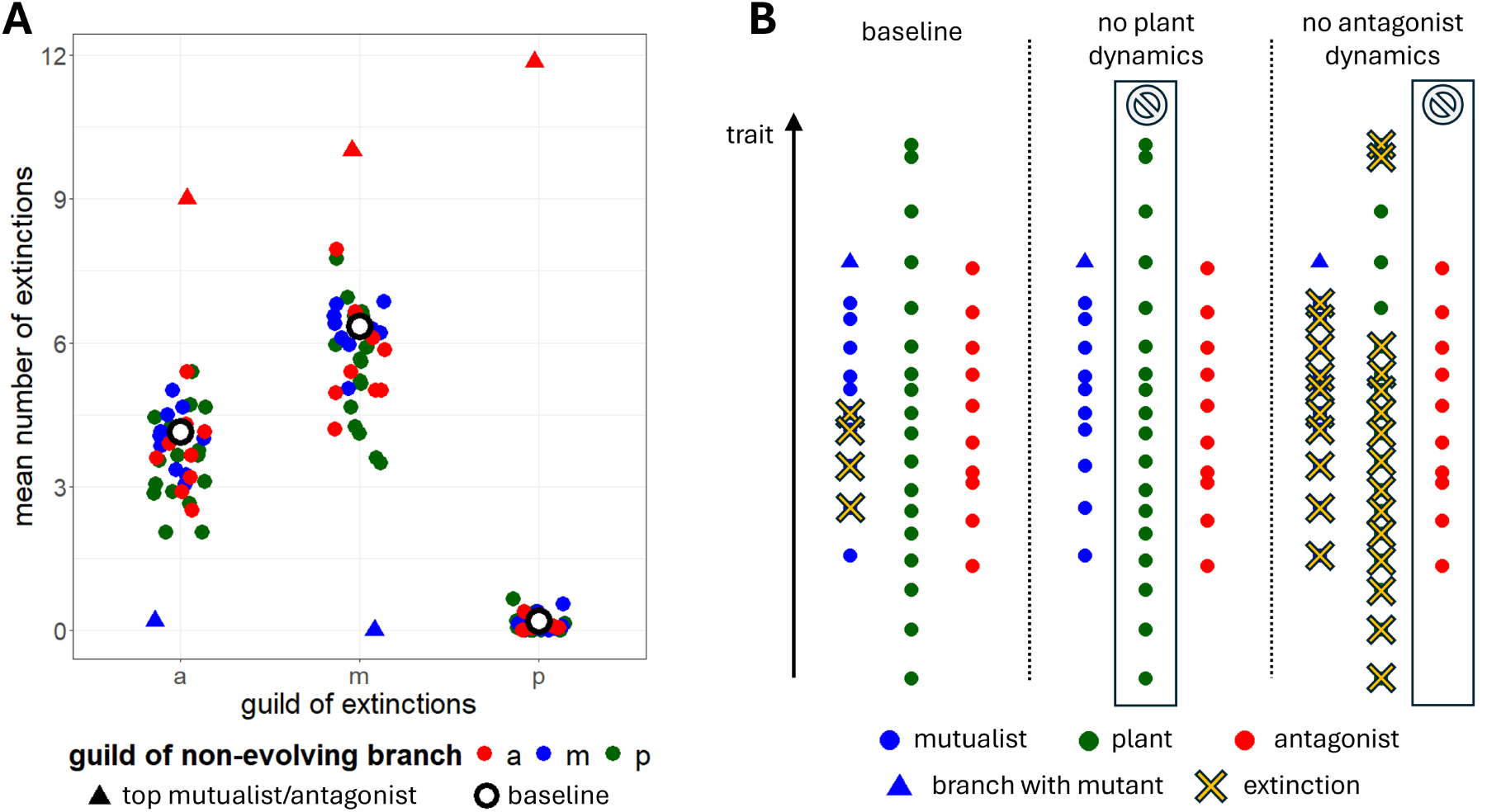
– Manipulative experiments. (A) Suppression of evolution of one branch at a time. Simulations are initialised with the state right before the first extinction avalanche in Figure 3 (black line there). Each point in the figure corresponds to a branch that is prohibited to evolve. The plot shows the number of extinctions in each of the guilds within 50 million time units, averaged over 20 replicates. The blue and red triangle correspond to the mutualist and antagonist branch with highest trait value (the only surviving branches in the original simulation run). Unfilled circles represent the baseline scenario with all branches evolving. (B) Manipulation of ecological dynamics after mutation, no further evolution. Simulations are initialised with the state right before the first extinction avalanche in Figure 3 (black line there), but with ecological dynamics at equilibrium and the next (mutualist) mutant added with initial density *N*_0_. The plot shows which branches go extinct after simulating the subsequent ecological dynamics (no evolution) for 100 000 time units in (i) the baseline scenario, (ii) a scenario with constant plant densities, (iii) a scenario with constant antagonist densities.

A second type of manipulative experiment demonstrates that the competitive exclusion of the other mutualists is fuelled by positive feedback from the mutualistic subnetwork (Fig. 4B). In this experiment, the mutant in the top mutualist branch causing the first extinctions in the original avalanche (extinction of 4 mutualist branches) is added to the system at ecological equilibrium. Subsequently, we tracked only the ecological dynamics without evolution, while switching off the density dynamics in single guilds. We find that extinctions can only occur when plant densities are allowed to change. This underlines that the mutualistic feedback is crucial for the extinctions to happen, since the additional indirect interactions increase effective competition within the mutualist guild (higher density of ‘winning’ mutualist *→* higher density of ‘winning’ plant(s) *→* lower density of other plants *→* lower density of other mutualists). Turning off only antagonist dynamics instead increases the number of extinctions, suggesting that feedbacks mediated by the antagonistic guild have a stabilising influence.

The extinction avalanche in antagonists in the original example quickly follows the avalanche in mutualists since the antagonist closest to the winning mutualist/plants has plant resources of higher density and outcompetes the other antagonists. After a period of an escape race between the remaining antagonist and mutualist, there may be an additional extinction avalanche in the plants (see Fig. 3B, after around 1.32 billion time units). Again, the ‘winning’ plants outcompete all other plant branches. The reason why the avalanche in plants does not happen as quickly as it does in the other guilds is that self-regulation is assumed much higher in plants (see Table 1), hence their densities are much less sensitive to feedbacks from the interaction network. The extinction avalanche in plants can only happen if the ‘winning’ mutualist reaches a particularly large lead over the next antagonist.

We were able to confirm our findings on the mechanisms of extinction avalanches using a broader set of example avalanches. First, the described manipulative experiments were repeated with five randomly chosen avalanches from simulations with the baseline parametrisation, yielding similar results (see Supplementary Material S5). Second, using 20 replicate runs with the baseline scenario, we could show that in 83.8% of all 74 avalanches in the mutualist guild, the lead of the mutualist over the antagonist (either at the top or bottom of the trait space) right before the avalanche was in the top percentile of leads since the last time that there were at most 4 mutualist branches (in 66.2% even maximal). We expect the remaining 16.2% of avalanches to follow the same pattern, but there are technical difficulties in determining the exact time of an avalanche if it happens to be preceded shortly beforehand by an extinction that is mechanistically unrelated (see Supplementary Material S2). This indicates that extinction avalanches usually occur after an unprecedented increase in the mutualist’s lead. The fact that the lead can also easily decrease and become negative (as observed for the bottom mutualist in Fig. 3E) furthermore suggests that the lead follows a random walk. Hence, although we can clearly identify conditions under which the network is more or less prone to extinction avalanches, their exact timing remains stochastic due to randomness in the mutational process.

The outlined mechanism explains the patterns in the prevalence of extinction avalanches observed in the previous section. If antagonism is stronger than mutualism or antagonists evolve fast compared to mutualists, the outermost antagonist branches tend to maintain a lead over the nearest mutualists, preventing extinction avalanches in the top left part of Figure 1A and B. Moreover, for higher general interaction strengths or higher general speed of antagonist and mutualist evolution, it takes less time until the lead of an outer mutualist over the next antagonist exceeds the threshold needed for triggering the avalanche. This explains why avalanche rate is higher in the top right part than in the bottom left part of Figure 1A and B.

Furthermore, the described mechanism requires trait-independent competition within the guilds to enable competitive exclusion across the entire trait space. This is in line with the substantial decrease of avalanche rates for high values of *α^P^*, extinction avalanches can then only occur if *α^M^* and *α^A^* are low (see Figure 2). The effect of *α^M^* and *α^A^* is more subtle for low to intermediate values of *α^P^*. Since a higher proportion of similarity-based competition leads to stronger competition-induced selection, *α^M^* and *α^A^* have a direct effect on the speed of mutualist and antagonist evolution. If *α^A^* is higher than *α^M^*, mutualists therefore do not reach plant branches ahead of antagonists. This explains the absence of extinction avalanches in the top left parts of Figure 2.

## Discussion

In this study, we find recurrent extinction avalanches that emerge from eco-evolutionary dynamics in a mutualistic-antagonistic interaction network without any environmental change. These extinction avalanches are triggered by a mutualist evolving to interact with plants that are not sufficiently controlled by antagonists. Subsequently, this mutualist, its plant partner(s) and the most successful antagonist outcompete a large proportion of the community. In general, we find that the balanced interplay of multiple interaction types is needed for avalanches to happen. Mutualism and trait-independent competition promote evolutionarily driven extinction events while antagonism and similarity-based competition are needed for the initial (and post-avalanche) diversification of the community.

The different interaction types contribute to the avalanche dynamics via a combination of eco-evolutionary mechanisms. First, trait-independent competition, reflecting reduced niche differences, generally increases the likelihood of competitive exclusion as soon as one type is better adapted to the biotic or abiotic environment than its competitors. This is coherent with previous theoretical investigations (Chesson, 2000; Leoz et al., 2026; Loeuille, 2019; Norberg et al., 2012; van Eldijk et al., 2020) and was recently confirmed in an evolution experiment involving bacteriophages under thermal stress (Greenrod et al., 2026). Second, in our model, mutualism further increases effective within-guild competition via additional indirect negative interactions. For example, a less abundant mutualist competitor suffers not only from direct competition with other mutualists but also from a low density of its plant partners due to their competition with other plants that interact with more abundant mutualists. Finally, contrary to competition and mutualism, antagonism stabilises the system via self-dampening feedbacks, enabling diversification in the first place, but once a mutualist overtakes an antagonist on their way to the unoccupied plant branches, the antagonist control is sufficiently weakened and the extinction avalanche is set in motion.

Our results contribute to a growing body of literature suggesting that competition and mutualism promote evolutionary murder (Leoz et al., 2026; Loeuille, 2019; Weinbach et al., 2022; Weyerer et al., 2023), while antagonism favours ‘indirect evolutionary rescue’, in which the evolution of one species saves its interaction partner from extinction (van Velzen, 2023; Yamamichi and Miner, 2015). However, the underlying mechanisms in these models vary considerably, and the results are not unequivocal. For instance, Shang et al. (2024) find that, in an antagonistic interaction, the prey may adapt in such a way as to escape a predator subjected to selective harvest, leading to evolutionary murder instead of indirect rescue. Also, in contrast to our study, Weinbach et al. (2022) and Weyerer et al. (2023) assume a trade-off between investing in mutualistic interactions and intrinsic growth, so that the mutualist, by investing less in the interaction, can murder its interaction partner – instead of its competitors, as in our case. We hypothesise that there is no general answer as to which type of biotic interaction induces which type of evolutionary outcome. This can depend on the presence and shape of trade-offs in the system as well as on the relative speed of evolution of interacting partners and potential environmental pressures.

The dynamics in our eco-evolutionary model share important features with the concept of ecological tipping points, with the difference that there is no external forcing involved. Instead, the gradual parameter change that induces the tipping emerges in a self-organised manner, that is, from the inherent system dynamics only. In our case, this parameter is the distribution of traits, in particular the lead of the outer mutualist over the outer antagonist, the increase in which may trigger the extinction avalanche. Our results are thus in line with previous theoretical work emphasising that evolution can trigger ecological tipping points (Ardichvili et al., 2023; Dakos et al., 2019), e.g., in the context of ‘evolutionary suicide’ (Dieckmann and Ferrière, 2004; Parvinen, 2005), or periodic evolutionary reversals (Dercole et al., 2002). Since the avalanches in our model are recurrent, the dynamics can also be seen as a higher-dimensional version of branching-extinction cycles as observed by Kisdi et al. (2002) and Dercole (2003), with ongoing changes in diversity and species composition rather than convergence towards an evolutionarily stable community (Edwards et al., 2018; Kremer and Klausmeier, 2017). Notably, Guill and Drossel (2008) and Allhoff et al. (2015) observed similar extinction avalanches in evolutionary food web models. They hypothesise an alternative underlying mechanism based on the evolved increase in diet specialisation of the species in their model. The resulting absence of generalist predators facilitates the invasion of mutants that face low predation pressure and can outcompete other species at their trophic level, with cascading bottom-up effects on higher trophic levels. According to this hypothesis, and in line with our findings, the lack of antagonistic control plays a crucial role in extinction avalanches.

The idea of self-organised tipping points resembles the concept of ‘self-organised criticality’ (SOC) which describes the tendency of a system to generically evolve towards a critical state in which it is susceptible to fluctuations whose magnitudes follow a power-law frequency distribution (Bak and Sneppen, 1993; Bak et al., 1987). However, we could not detect a power law in the size distribution of avalanches. This adds to the mixed evidence for such a power law from paleobiological data (Newman and Palmer, 2003) as well as from evolutionary network models (Allhoff et al., 2015; Drossel et al., 2001; Guill and Drossel, 2008; Loeuille and Loreau, 2005), which questions the applicability of SOC to eco-evolutionary community dynamics.

Our results raise the question of how frequently evolution-induced species extinctions occur in nature (Rankin and López-Sepulcre, 2005; Webb, 2003). Giving an answer to this question is notoriously challenging for a number of reasons. First, detecting extinctions is inherently difficult, e.g., when estimating extinction rates from molecular phylogenies (Louca and Pennell, 2021; Rabosky, 2010), or when attempting to infer local extirpations from observational data (Simon et al., 2026). Second, attributing an extinction to a certain cause is most often impossible, since-due to the absence of experimental data studies can usually only establish correlation rather than causation (Gurevitch and Padilla, 2004). Subject to these limitations, species extinctions are most often regarded as the result of external disturbances or purely ecological dynamics within communities (Bond and Grasby, 2017; Sodhi et al., 2009). In contrast, our findings suggest that extinctions may be linked to a history of coevolution, particularly between the ‘winning’ mutualist and its plant partners. The signal of this coevolution could possibly be identified using phenotypic, genomic and/or phylogenetic data (Anderson, 2015). Moreover, recent advances in experimental evolution on the community scale-potentially including different types of biotic interactions-offer hope that, in future, the predictions of eco-evolutionary models such as ours can be tested more accurately (Montbel and Hrcek, 2026).

Our modelling approach is inevitably subject to certain limitations. All biotic interactions of an organism in our model are mediated by a single trait, an instance of ‘ecological pleiotropy’ (*sensu* Strauss and Irwin, 2004). This need not be the case in nature; for instance, the plant traits that mainly govern the interaction with pollinators (e.g., flower shape) may differ from those that influence herbivory (e.g., defence compounds), and may evolve independently from the latter. It has been shown that the absence of ecological pleiotropy decouples diversification patterns in mutualistic and antagonistic subnetworks and reduces the potential for extinction avalanches (Jäger et al., 2026c). Moreover, while the trait matching rule that determines interaction strengths in our model is widely used in theoretical literature (Allhoff et al., 2015; Doebeli and Dieckmann, 2000; Kiester et al., 1984; Loeuille and Loreau, 2005; Yacine and Loeuille, 2024), empirical support is mixed (Bartomeus et al., 2016; Dormann et al., 2017; Schurr et al., 2025). Any change to this assumption would fundamentally alter the structure of the emerging interaction networks and feedback loops in the model, with implications for extinction avalanches that have yet to be investigated.

Whilst the large-scale mass extinctions in Earth history are undoubtedly often linked to environmental changes, we have shown that, within isolated local communities, evolution can trigger the collapse of large parts of ecological networks without any external forcing. To the best of our knowledge, our study is the first to provide a mechanism underlying such extinction avalanches. In particular, the combination of different interaction types opens up new avenues of evolutionary murder, potentially claiming many victims simultaneously. Although this is extremely difficult to prove empirically, future research should be aware of (co)evolution as a potential driving force behind extinction events.

## Acknowledgements

FJ would like to thank Toni Klauschies and Frank Schurr for insightful discussions. The authors furthermore acknowledge support by the state of Baden-Württemberg through bwHPC.

## Data, script, code, and supplementary information availability

All data, script and code for this work are available online (https://zenodo.org/records/22874553); Jäger et al., 2026a. Supplementary information is available online (https://zenodo.org/records/22877472; Jäger et al., 2026b).

## References

1. Allhoff KT, Ritterskamp D, Rall BC, Drossel B, Guill C (2015). Evolutionary Food Web Model Based on Body Masses Gives Realistic Networks with Permanent Species Turnover. Scientific Reports 5, 10955. 10.1038/srep10955. (Visited on 07/30/2026).

2. Anderson B (2015). Coevolution in Mutualisms. In: Mutualism. Ed. by Judith L. Bronstein. Oxford: Oxford University Press, pp. 107–130. 10.1093/acprof:oso/9780199675654.003.0007. (Visited on 03/19/2024).

3. Apanius V, Penn D, Slev PR, Ruff LR, Potts WK (1997). The Nature of Selection on the Major Histocompatibility Complex. Critical Reviews in Immunology 17, 179–224. 10.1615/CritRevImmunol.v17.i2.40. (Visited on 08/27/2026).

4. Ardichvili AN, Loeuille N, Dakos V (2023). Evolutionary Emergence of Alternative Stable States in Shallow Lakes. Ecology Letters 26, 692–705. 10.1111/ele.14180. (Visited on 08/27/2026).

5. Bak P, Sneppen K (1993). Punctuated Equilibrium and Criticality in a Simple Model of Evolution. Physical Review Letters 71, 4083–4086. 10.1103/PhysRevLett.71.4083. (Visited on 01/16/2026).

6. Bak P, Tang C, Wiesenfeld K (1987). Self-Organized Criticality: An Explanation of the 1/f Noise. Physical Review Letters 59, 381–384. 10.1103/PhysRevLett.59.381. (Visited on 07/30/2026).

7. Bartomeus I, Gravel D, Tylianakis JM, Aizen MA, Dickie IA, Bernard-Verdier M (2016). A Common Framework for Identifying Linkage Rules across Different Types of Interactions. Functional Ecology 30, 1894–1903. 10.1111/1365-2435.12666. (Visited on 10/21/2024).

8. Bellard C, Genovesi P, Jeschke JM (2016). Global Patterns in Threats to Vertebrates by Biological Invasions. Proceedings of the Royal Society B: Biological Sciences 283, 20152454. 10.1098/rspb.2015.2454. (Visited on 07/14/2026).

9. Bond DPG, Grasby SE (2017). On the Causes of Mass Extinctions. Palaeogeography, Palaeoclimatology, Palaeoecology. Mass Extinction Causality: Records of Anoxia, Acidification, and Global Warming during Earth’s Greatest Crises 478, 3–29. 10.1016/j.palaeo.2016.11.005. (Visited on 06/02/2026).

10. Brook BW, Sodhi NS, Ng PKL (2003). Catastrophic Extinctions Follow Deforestation in Singapore. Nature 424, 420–423. 10.1038/nature01795. (Visited on 07/15/2026).

11. Burbidge AA, Manly BFJ (2002). Mammal Extinctions on Australian Islands: Causes and Conservation Implications. Journal of Biogeography 29, 465–473. 10.1046/j.1365-2699.2002.00699.x. (Visited on 07/14/2026).

12. Chesson P (2000). Mechanisms of Maintenance of Species Diversity. Annual Review of Ecology, Evolution, and Systematics 31, 343–366. 10.1146/annurev.ecolsys.31.1.343. (Visited on 07/14/2026).

13. Dakos V, Bascompte J (2014). Critical Slowing down as Early Warning for the Onset of Collapse in Mutualistic Communities. Proceedings of the National Academy of Sciences 111, 17546– 17551. 10.1073/pnas.1406326111. (Visited on 07/30/2026).

14. Dakos V, Matthews B, Hendry AP, Levine J, Loeuille N, Norberg J, Nosil P, Scheffer M, De Meester L (2019). Ecosystem Tipping Points in an Evolving World. Nature Ecology & Evolution 3, 355–362. 10.1038/s41559-019-0797-2. (Visited on 01/18/2024).

15. Dercole F (2003). Remarks on Branching-Extinction Evolutionary Cycles. Journal of Mathematical Biology 47, 569–580. 10.1007/s00285-003-0236-4. (Visited on 07/29/2026).

16. Dercole F, Ferriere R, Rinaldi S (2002). Ecological Bistability and Evolutionary Reversals under Asymmetrical Competition. Evolution 56, 1081–1090. 10.1111/j.0014-3820.2002.tb01422.x. (Visited on 07/29/2026).

17. Dieckmann U, Ferrière R (2004). Adaptive Dynamics and Evolving Biodiversity. In: Evolutionary Conservation Biology. Ed. by Denis Couvet, Régis Ferrière, and Ulf Dieckmann. Cambridge Studies in Adaptive Dynamics. Cambridge: Cambridge University Press, pp. 188–224. 10.1017/CBO9780511542022.015. (Visited on 07/30/2026).

18. Doebeli M, Dieckmann U (2000). Evolutionary Branching and Sympatric Speciation Caused by Different Types of Ecological Interactions. The American Naturalist 156, S77–S101. 10.1086/303417. (Visited on 02/19/2025).

19. Donohue I, Petchey OL, Kéfi S, Génin A, Jackson AL, Yang Q, O’Connor NE (2017). Loss of Predator Species, Not Intermediate Consumers, Triggers Rapid and Dramatic Extinction Cascades. Global Change Biology 23, 2962–2972. 10.1111/gcb.13703. (Visited on 07/15/2026).

20. Dormann CF, Fründ J, Schaefer HM (2017). Identifying Causes of Patterns in Ecological Networks: Opportunities and Limitations. Annual Review of Ecology, Evolution, and Systematics 48, 559–584. 10.1146/annurev-ecolsys-110316-022928. (Visited on 11/29/2024).

21. Drossel B, Higgs PG, McKane AJ (2001). The Influence of Predator–Prey Population Dynamics on the Long-Term Evolution of Food Web Structure. Journal of Theoretical Biology 208, 91–107. 10.1006/jtbi.2000.2203. (Visited on 07/30/2026).

22. Edwards KF, Kremer CT, Miller ET, Osmond MM, Litchman E, Klausmeier CA (2018). Evolutionarily Stable Communities: A Framework for Understanding the Role of Trait Evolution in the Maintenance of Diversity. Ecology Letters 21, 1853–1868. 10.1111/ele.13142. (Visited on 07/30/2026).

23. Ester M, Kriegel HP, Sander J, Xu X (1996). A Density-Based Algorithm for Discovering Clusters in Large Spatial Databases with Noise. In: Proceedings of the Second International Conference on Knowledge Discovery and Data Mining. KDD’96. Portland, Oregon: AAAI Press, pp. 226–231. (Visited on 02/18/2026).

24. Estes JA, Burdin A, Doak DF (2016). Sea Otters, Kelp Forests, and the Extinction of Steller’s Sea Cow. Proceedings of the National Academy of Sciences 113, 880–885. 10.1073/pnas.1502552112. (Visited on 07/15/2026).

25. Fiegna F, Velicer GJ (2003). Competitive Fates of Bacterial Social Parasites: Persistence and Self– Induced Extinction of Myxococcus Xanthus Cheaters. Proceedings of the Royal Society B: Biological Sciences 270, 1527–1534. 10.1098/rspb.2003.2387. (Visited on 07/14/2026).

26. Fontaine C, Guimarães Jr PR, Kéfi S, Loeuille N, Memmott J, van der Putten WH, van Veen FJF, Thébault E (2011). The Ecological and Evolutionary Implications of Merging Different Types of Networks. Ecology Letters 14, 1170–1181. 10.1111/j.1461-0248.2011.01688.x. (Visited on 02/28/2023).

27. Futuyma DJ, Agrawal AA (2009). Macroevolution and the Biological Diversity of Plants and Herbivores. Proceedings of the National Academy of Sciences 106, 18054–18061. 10.1073/pnas.0904106106. (Visited on 01/30/2026).

28. Greenrod STE, Cazares D, Ślesak W, Hector TE, MacLean RC, King KC (2026). Rapid Adaptation Accelerates Competitive Suppression in a Parasite Community. The ISME Journal 20, wrag114. 10.1093/ismejo/wrag114. (Visited on 07/14/2026).

29. Guill C, Drossel B (2008). Emergence of Complexity in Evolving Niche-Model Food Webs. Journal of Theoretical Biology 251, 108–120. 10.1016/j.jtbi.2007.11.017. (Visited on 01/16/2026).

30. Gurevitch J, Padilla DK (2004). Are Invasive Species a Major Cause of Extinctions? Trends in Ecology & Evolution 19, 470–474. 10.1016/j.tree.2004.07.005. (Visited on 09/09/2026).

31. Hackett TD, Sauve AMC, Davies N, Montoya D, Tylianakis JM, Memmott J (2019). Reshaping Our Understanding of Species’ Roles in Landscape-Scale Networks. Ecology Letters 22, 1367– 1377. 10.1111/ele.13292. (Visited on 10/02/2025).

32. Hallam A, Wignall PB (1997). Mass Extinctions and Their Aftermath. Oxford: Oxford University Press. 10.1093/oso/9780198549178.001.0001. (Visited on 06/16/2026).

33. Jäger F, Guill C, Loeuille N, Yacine Y, Allhoff KT (2026a). Eco-evolutionary dynamics create extinction avalanches in mutualistic-antagonistic networks-Data and Code. Zenodo. 10.5281/zenodo.22874552.

34. Jäger F, Guill C, Loeuille N, Yacine Y, Allhoff KT (2026b). Eco-evolutionary dynamics create extinction avalanches in mutualistic-antagonistic networks-Supplementary Material. Zenodo. 10.5281/zenodo.22877471.

35. Jäger F, Loeuille N, Yacine Y, Allhoff KT (2026c). Between Friends and Foes: Evolutionary Diversification in Mutualistic-Antagonistic Networks. bioRxiv (Preprint). 10.64898/2026.03.16.712075. (Visited on 07/24/2026).

36. Kéfi S, Berlow EL, Wieters EA, Joppa LN, Wood SA, Brose U, Navarrete SA (2015). Network Structure beyond Food Webs: Mapping Non-Trophic and Trophic Interactions on Chilean Rocky Shores. Ecology 96, 291–303. 10.1890/13-1424.1. (Visited on 10/14/2024).

37. Kiester AR, Lande R, Schemske DW (1984). Models of Coevolution and Speciation in Plants and Their Pollinators. The American Naturalist 124, 220–243. 10.1086/284265. JSTOR:2461492. (Visited on 05/22/2025).

38. Kisdi É, Jacobs FJA, Geritz SAH (2002). Red Queen Evolution by Cycles of Evolutionary Branching and Extinction. Selection 2, 161–176. 10.1556/select.2.2001.1-2.12. (Visited on 07/29/2026).

39. Kremer CT, Klausmeier CA (2017). Species Packing in Eco-Evolutionary Models of Seasonally Fluctuating Environments. Ecology Letters 20, 1158–1168. 10.1111/ele.12813. (Visited on 07/30/2026).

40. Leoz S, Lutscher F, Allhoff KT, Govaert L (2026). Eco-Evolutionary Dynamics in Competitive Systems: Rescue and Murder. bioRxiv (Preprint). 10.64898/2026.01.14.699340. (Visited on 07/14/2026).

41. Lever JJ, van Nes EH, Scheffer M, Bascompte J (2014). The Sudden Collapse of Pollinator Communities. Ecology Letters 17, 350–359. 10.1111/ele.12236. (Visited on 07/30/2026).

42. Loeuille N (2010). Influence of Evolution on the Stability of Ecological Communities. Ecology Letters 13, 1536–1545. 10.1111/j.1461-0248.2010.01545.x. (Visited on 11/14/2024).

43. Loeuille N (2019). Eco-Evolutionary Dynamics in a Disturbed World: Implications for the Maintenance of Ecological Networks [Version 1; Peer Review: 2 Approved]. F1000Research 8, 97. 10.12688/f1000research.15629.1. (Visited on 10/21/2024).

44. Loeuille N, Loreau M (2005). Evolutionary Emergence of Size-Structured Food Webs. Proceedings of the National Academy of Sciences 102, 5761–5766. 10.1073/pnas.0408424102. (Visited on 02/28/2023).

45. Louca S, Pennell MW (2021). Why Extinction Estimates from Extant Phylogenies Are so Often Zero. Current Biology 31, 3168–3173.e4. 10.1016/j.cub.2021.04.066. (Visited on 09/10/2026).

46. McCann KS (2000). The Diversity–Stability Debate. Nature 405, 228–233. 10.1038/35012234. (Visited on 08/27/2026).

47. Melián CJ, Bascompte J, Jordano P, Krivan V (2009). Diversity in a Complex Ecological Network with Two Interaction Types. Oikos 118, 122–130. 10.1111/j.1600-0706.2008.16751.x. (Visited on 10/17/2024).

48. Miller AH, Stroud JT, Losos JB (2023). The Ecology and Evolution of Key Innovations. Trends in Ecology & Evolution 38, 122–131. 10.1016/j.tree.2022.09.005. (Visited on 07/14/2026).

49. Miller RR, Williams JD, Williams JE (1989). Extinctions of North American Fishes during the Past Century. Fisheries 14, 22–38. 10.1577/1548-8446(1989)014<0022:EONAFD>2.0.CO;2. (Visited on 07/14/2026).

50. Montbel V, Hrcek J (2026). Experimental Evolution in Communities: Beyond Pairwise Interactions. Evolution 80, 889–901. 10.1093/evolut/qpag029. (Visited on 09/09/2026).

51. Morrison BML, Brosi BJ, Dirzo R (2020). Agricultural Intensification Drives Changes in Hybrid Network Robustness by Modifying Network Structure. Ecology Letters 23, 359–369. 10.1111/ele.13440. (Visited on 08/22/2025).

52. Newell ND (1967). Revolutions in the History of Life. In: Uniformity and Simplicity: A Symposium on the Principle of the Uniformity of Nature. Ed. by Claude C. Albritton Jr. Vol. 89. New York: Geological Society of America, p. 0. 10.1130/SPE89-p63. (Visited on 06/16/2026).

53. Newman MEJ, Palmer RG (2003). Modeling Extinction. Oxford: Oxford University Press. 10.1093/oso/9780195159455.001.0001. (Visited on 07/30/2026).

54. Norberg J, Urban MC, Vellend M, Klausmeier CA, Loeuille N (2012). Eco-Evolutionary Responses of Biodiversity to Climate Change. Nature Climate Change 2, 747–751. 10.1038/nclimate1588. (Visited on 07/31/2026).

55. Olsen EM, Heino M, Lilly GR, Morgan MJ, Brattey J, Ernande B, Dieckmann U (2004). Maturation Trends Indicative of Rapid Evolution Preceded the Collapse of Northern Cod. Nature 428, 932–935. 10.1038/nature02430. (Visited on 07/14/2026).

56. Paine RT (1966). Food Web Complexity and Species Diversity. The American Naturalist 100, 65–75. 10.1086/282400. (Visited on 07/15/2026).

57. Parvinen K (2005). Evolutionary Suicide. Acta Biotheoretica 53, 241–264. 10.1007/s10441-005-2531-5. (Visited on 07/14/2026).

58. Pocock MJO, Evans DM, Memmott J (2012). The Robustness and Restoration of a Network of Ecological Networks. Science 335, 973–977. 10.1126/science.1214915. (Visited on 10/16/2024).

59. Rabosky DL (2010). Extinction Rates Should Not Be Estimated from Molecular Phylogenies. Evolution 64, 1816–1824. 10.1111/j.1558-5646.2009.00926.x. (Visited on 09/10/2026).

60. Rankin DJ, López-Sepulcre A (2005). Can Adaptation Lead to Extinction? Oikos 111, 616–619. 10.1111/j.1600-0706.2005.14541.x. (Visited on 09/16/2026).

61. Raup DM (1986). Biological Extinction in Earth History. Science 231, 1528–1533. 10.1126/science.11542058. (Visited on 01/19/2026).

62. Rozen DE, Lenski RE (2000). Long-Term Experimental Evolution in Escherichia Coli. VIII. Dynamics of a Balanced Polymorphism. The American Naturalist 155, 24–35. 10.1086/303299.

63. Sanders D, Sutter L, van Veen FJF (2013). The Loss of Indirect Interactions Leads to Cascading Extinctions of Carnivores. Ecology Letters 16, 664–669. 10.1111/ele.12096. (Visited on 07/15/2026).

64. Schurr F, Jäger F, Cappellari-Rabeling S, Grass I, Pagel J, Palmer M, Petschenka G, Razanajatovo M, Schlüter PM, Schweiger AH, Sheppard CS, Steppuhn A, Allhoff KT (2025). Fitness Landscapes of Biotic Interactions Shape the Ecological and Evolutionary Dynamics of Biodiversity. EcoEvoRxiv (Preprint). 10.32942/X2BQ0Z. (Visited on 02/25/2026).

65. Shang Y, Kasada M, Kondoh M (2024). Rescue or Murder? The Effect of Prey Adaptation to the Predator Subjected to Fisheries. Ecology and Evolution 14, e70336. 10.1002/ece3.70336. (Visited on 07/14/2026).

66. Simon ADF, Basman A, Martin RA, Robinson C, Cronk Q (2026). Detecting Extirpation: A Localized Approach to a Global Problem. Plants, People, Planet 8, 1160–1174. 10.1002/ppp3.70130. (Visited on 09/09/2026).

67. Sodhi NS, Brook BW, Bradshaw CJA (2009). Causes and Consequences of Species Extinctions. In: The Princeton Guide to Ecology. Ed. by Simon A. Levin. Princeton: Princeton University Press. Chap. The Princeton Guide to Ecology, pp. 514–520. 10.1515/9781400833023.514. (Visited on 09/09/2026).

68. Strauss SY, Irwin RE (2004). Ecological and Evolutionary Consequences of Multispecies Plant-Animal Interactions. Annual Review of Ecology, Evolution, and Systematics 35, 435–466. 10.1146/annurev.ecolsys.35.112202.130215. (Visited on 12/03/2024).

69. van Eldijk TJB, Bisschop K, Etienne RS (2020). Uniting Community Ecology and Evolutionary Rescue Theory: Community-wide Rescue Leads to a Rapid Loss of Rare Species. Frontiers in Ecology and Evolution 8, 552268. 10.3389/fevo.2020.552268. (Visited on 07/31/2026).

70. van Velzen E (2023). High Importance of Indirect Evolutionary Rescue in a Small Food Web. Ecology Letters 26, 2110–2121. 10.1111/ele.14321. (Visited on 06/02/2026).

71. Webb C (2003). A Complete Classification of Darwinian Extinction in Ecological Interactions. The American Naturalist 161, 181–205. 10.1086/345858. (Visited on 09/16/2026).

72. Weber MG, Agrawal AA (2014). Defense Mutualisms Enhance Plant Diversification. Proceedings of the National Academy of Sciences 111, 16442–16447. 10.1073/pnas.1413253111. (Visited on 06/23/2025).

73. Weinbach A, Loeuille N, Rohr RP (2022). Eco-Evolutionary Dynamics Further Weakens Mutualistic Interaction and Coexistence under Population Decline. Evolutionary Ecology 36, 373–387. 10.1007/s10682-022-10176-7. (Visited on 07/14/2026).

74. Weyerer F, Weinbach A, Zarfl C, Allhoff KT (2023). Eco-Evolutionary Dynamics in Two-Species Mutualistic Systems: One-Sided Population Decline Triggers Joint Interaction Disinvestment. Evolutionary Ecology 37, 981–999. 10.1007/s10682-023-10264-2. (Visited on 09/10/2024).

75. Yacine Y, Loeuille N (2024). Attracting pollinators vs escaping herbivores: eco-evolutionary dynamics of plants confronted with an ecological trade-off. Peer Community Journal 4, e60. 10.24072/pcjournal.433. (Visited on 11/27/2024).

76. Yamamichi M, Miner BE (2015). Indirect Evolutionary Rescue: Prey Adapts, Predator Avoids Extinction. Evolutionary Applications 8, 787–795. 10.1111/eva.12295. (Visited on 07/14/2026).

