## Supplementary Material for "Eco-evolutionary dynamics create extinction avalanches in mutualistic-antagonistic networks"

Felix Jäger[1,2], Christian Guill[3], Nicolas Loeuille[4], Youssef Yacine[5], and Korinna T. Allhoff[1,2,6,7]

DOI not yet assigned

### Abstract

Evolution has given rise to the diversity of life on Earth, but it can also cause species extinctions. While major extinction events are usually associated with external drivers, in this study, we show mechanistically how evolution can trigger abrupt extinction avalanches in ecological networks without any environmental change. In particular, we investigate an eco-evolutionary simulation model of a tripartite mutualistic-antagonistic network, such as a plant-pollinator-herbivore network, with additional intraguild competition. The prevalence of self-organised extinction avalanches depends on a subtle balance of the different interaction types, where antagonism and similarity-based competition are needed for initial diversification while mutualism and trait-independent competition foster evolutionarily driven extinction events. Extinction avalanches are triggered by a mutualist evolving to interact with plants that are not subject to sufficient antagonistic control. Subsequently, the mutualist, its plant partners and their main antagonist potentially outcompete the rest of the community. Our findings point out that evolutionary murder may claim many victims simultaneously and highlight the need to take different interaction types into account to gain a comprehensive understanding of the links between evolution and biodiversity in ecological communities.

[1]Department of Eco-Evolutionary Modelling (190m), University of Hohenheim, Stuttgart, Germany, [2]Computational Science Hub (CSH), University of Hohenheim, Stuttgart, Germany, [3]Institute for Chemistry and Biology of the Marine Environment (ICBM), Carl von Ossietzky University Oldenburg, Oldenburg, Germany, [4]Sorbonne Université, Université Paris Cité, Univ Paris Est Créteil, CNRS, IRD, INRAE, Institut d’Écologie et des Sciences de l’Environnement de Paris (iEES-Paris), Paris, France, [5]MARBEC, Université de Montpellier, CNRS, Ifremer, IRD, Montpellier, France, [6]KomBioTa - Center for Biodiversity and Integrative Taxonomy, University of Hohenheim & State Museum of Natural History, Stuttgart, Germany, [7]Cluster of Excellence GreenRobust, University of Hohenheim, Stuttgart, Germany

**Correspondence**


### S1. Robustness to quantitative changes in avalanche criterion

We repeated the analyses of extinction avalanche rates while varying the parameters in the extinction avalanche criterion. In particular, we varied the maximum time window in which the avalanche can occur (100 million time units in the main text) and the minimum proportion of branches that has to be lost within that period (0.5 in the in the main text). Figure S1 shows the results for varying antagonism/mutualism strength and varying relative evolutionary speeds. We observe that changing the maximum time window has virtually no effect while a higher minimum relative loss results in fewer recorded extinction avalanches. However, the qualitative patterns do not change. This was also confirmed when varying the proportions of similarity-based competition $\alpha^A$, $\alpha^M$, $\alpha^P$ (not shown).

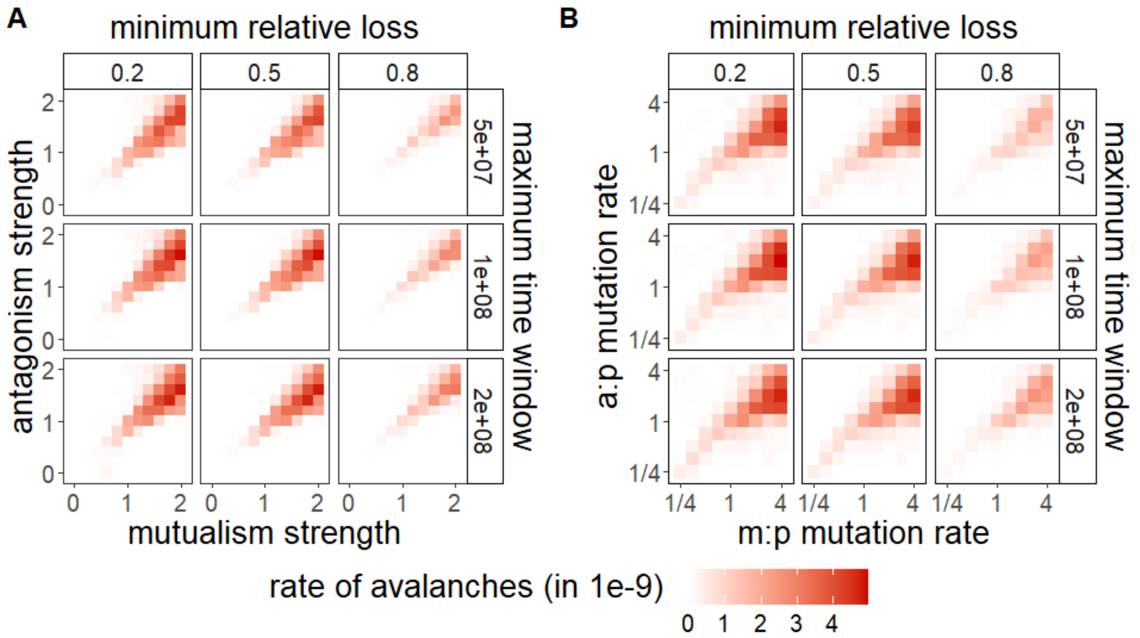

**Figure S1 – Rate of extinction avalanches under varying extinction avalanche criteria.** Plots show the rate of extinction avalanches, measured as the total number of extinction avalanches in any of the guilds over total simulated time, using 5 replicates for each parameter combination. The rate is computed for varying minimum relative loss and maximum time window in the extinction avalanche criterion, as well as (A) varying maximum strength of mutualism and antagonism ($a_0^{mut}$ and $a_0^{ant}$), and (B) varying relative mutualist and antagonist mutation rates ($\mu_M : \mu_P$ and $\mu_A : \mu_P$; $\mu_P = 2 \cdot 10^{-6}$). All other parameters were chosen according to Table 1 from the main text.

### S2. Calculation of extinction avalanche metrics

In this section, we explain in more detail the methods used to determine the number of extinction avalanches, their size, the order of extinction avalanches in the different guilds and the precise starting time of an extinction avalanche.

In general, an extinction avalanche was defined as the loss of more than 50% of the branches of a guild, but at least 4 branches, within at most 100 million time units. We call a time point $t$ 'critical' if there is an extinction avalanche happening within the next 100 million time units, i.e., if there are $t'$ and $t''$ with $t \leq t' < t''$ and $t'' - t \leq 10^8$, such that $n(t') - n(t'') > 0.5n(t')$ and $n(t') - n(t'') \geq 4$, where $n(t)$ is the number of branches present at time $t$. The number of extinction avalanches in a simulation run was defined as the number of sequences of consecutive critical time points (see Figure S2). The size of an avalanche is the maximum number of branches lost within 100 million time units after a critical time point in such a sequence.

In order to determine the order of extinction avalanches in different guilds in a simulation run, we compared the first critical time point in each guild. This was repeated for 20 replicate simulation runs with the baseline parametrisation. We found that in all replicate runs, extinction avalanches happened first in mutualists, then in antagonists and finally, if at all, in plants. Note that this method accounts only for the order of the first extinction avalanches in each simulation run. Extending this analysis to later extinction avalanches would require to define a rule to group extinction avalanches in different guilds that belong together. This is not readily possible in a canonical manner.

#### Analysis of the lead of the outermost mutualist branch at the onset of an extinction avalanche

Determining the exact start of an extinction avalanche is a non-trivial task. We used the following method (see Fig. S2). First, we recorded for each critical point in time $t$ the time until the extinction avalanche is over, which is the minimal possible value of $t'' - t$ such that there is a $t'$ with $t \leq t' < t''$, $n(t') - n(t'') > 0.5n(t')$ and $n(t') - n(t'') \geq 4$, where $n(t)$ is the number of branches present at time $t$. The beginning of an avalanche $t_{start}$ was now defined as the first point within a sequence of consecutive critical points in time where the 'time until extinction avalanche is over' stops decreasing. Note that this definition (like any alternative definition) is not ideal. In our simulation runs, for example, it occasionally happens that single branches are lost outside of extinction avalanches, for different reasons. If such a single branch is lost and, shortly afterwards, an extinction avalanche occurs according to the mechanism described in the main text, it may happen that the onset of the extinction avalanche is mistakenly associated with the prior loss of the single branch (cf. Fig. S2B).

The lead of the outermost mutualist over the outermost antagonist was defined as

$$\max(q^M_{top} - q^A_{top}, q^A_{bot} - q^M_{bot}),$$

where $q^M_{top}$ ($q^A_{top}$) is the maximum mutualist (antagonist) trait value and $q^M_{bot}$ ($q^A_{bot}$) the minimum mutualist (antagonist) trait value. For each mutualist extinction avalanche in 20 replicate runs with the baseline parametrisation (74 avalanches in total), we calculated this lead for each time point after the last time there were at most 4 mutualist branches (to allow only time points where

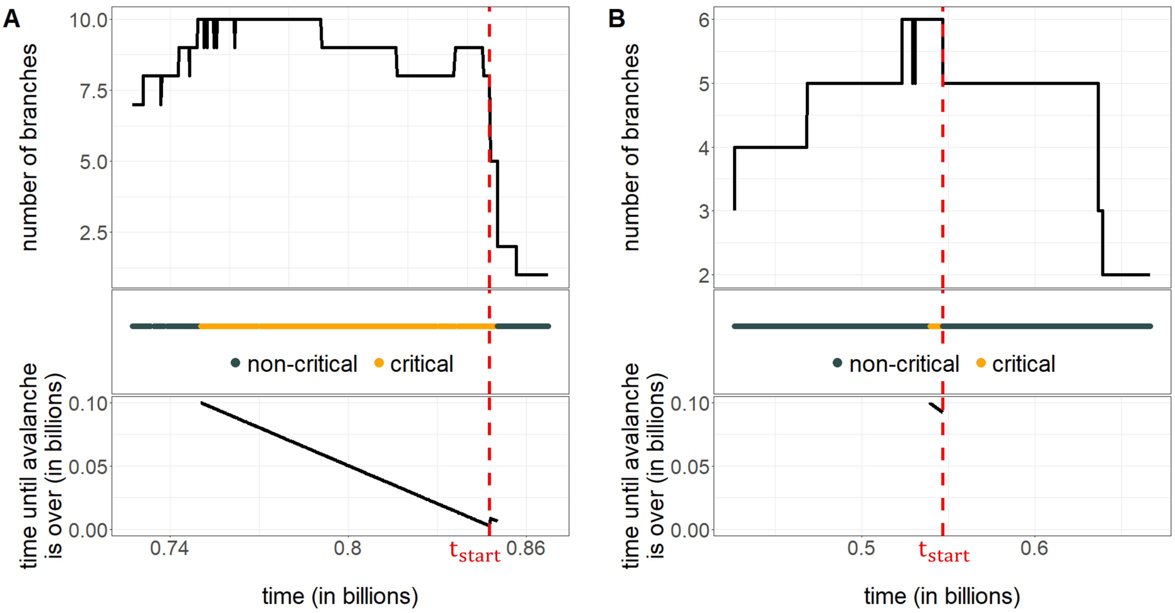

**Figure S2 – Determining the start of an avalanche**. Extinction avalanches are detected based on the number of branches in a guild over time (top panels, the two examples A and B are taken from extinction avalanches in the mutualist guild using the baseline parametrisation). Points in time are marked as 'critical' if there is an avalanche happening within the next 100 million time units. One extinction avalanche then corresponds to one sequence of consecutive critical time points (middle panel). For each critical time point, the 'time until avalanche is over' is calculated as the minimal time within which an extinction avalanche has happened (bottom panel). The start of an avalanche is defined as the first time point at which the 'time until avalanche is over' stops decreasing (dotted red lines). (A) Example of a successful identification of the start of an avalanche. (B) Example of a potential mis-identification of the start of an avalanche. The first extinction happens significantly earlier than the bulk of the extinctions, suggesting that it might be mechanistically unrelated.

avalanches are possible) until the computed beginning of the avalanche $t_{start}$. Subsequently, we computed the percentile of the lead at $t_{start}$ within all the leads in this period. In 83.8% of the extinction avalanches, the lead right before the avalanche was in the top percentile (in 66.2% even maximal). In 11 of the 12 avalanches where the lead was not in the top percentile, we could confirm that the bulk of extinctions happened significantly later than the calculated beginning of the extinction avalanche, suggesting a 'wrong' identification of the beginning of the extinction avalanche due to the issue described in the previous paragraph (as in Fig. S2B). In the remaining case, the avalanche seems to be triggered by the outermost mutualist branch overtaking the second-outermost antagonist branch while the outermost antagonist branch is far away in the trait space.

### S3. Diversification patterns under varying parameter combinations

Sufficient initial diversification is a necessary prerequisite for extinction avalanches to happen. In this section, we report results on diversification for all parameter combinations analysed in the main text. As a proxy for diversification, we use the maximum number of branches in each guild within a simulation run, averaged over 5 replicates.

First, we varied the maximum strength of mutualism and antagonism (Fig. S3A). We observe low diversification in all guilds if mutualism is stronger than antagonism, restricting the potential for extinction avalanches. In this case, stabilising selection exerted by the mutualistic partner prevents diversification in the plants and hence in all other guilds. If, on the other hand, antagonism strength exceeds mutualism strength, disruptive selection induced by antagonism and competition outweighs the stabilising selection and leads to diversification; first in the plant guild, and subsequently also in the other guilds. A similar pattern can be observed when varying the relative mutation rates of mutualists and antagonists (Fig. S3B). Faster evolution of mutualists or antagonists leads to better adaptation to plants and hence higher relative importance of mutualism- vs. antagonism-induced selection, with the described consequences for diversification patterns. Figure S3 also shows that plants tend to diversify more than the other guilds in our model, which can be explained by the fact that both mutualists and antagonists rely on plants for their survival whereas the reverse is not the case.

Finally, we varied the proportions of similarity-based competition (Fig. S4). We find that without similarity-based competition in the plants ($\alpha^P = 0$) diversification remains mostly limited. For low values of $\alpha^P$ ($0.1$ or $0.2$), diversification is favoured if $\alpha^A$ is greater than $\alpha^M$, meaning that there is more similarity-based competition in antagonists than in mutualists. This pattern follows from the observations made for relative speed of evolution since a higher proportion of similarity-based competition implies stronger competition-induced selection and therefore a higher speed of evolution. If competition in plants is mainly similarity-based (high $\alpha^P$), there is plenty of diversification in all guilds.

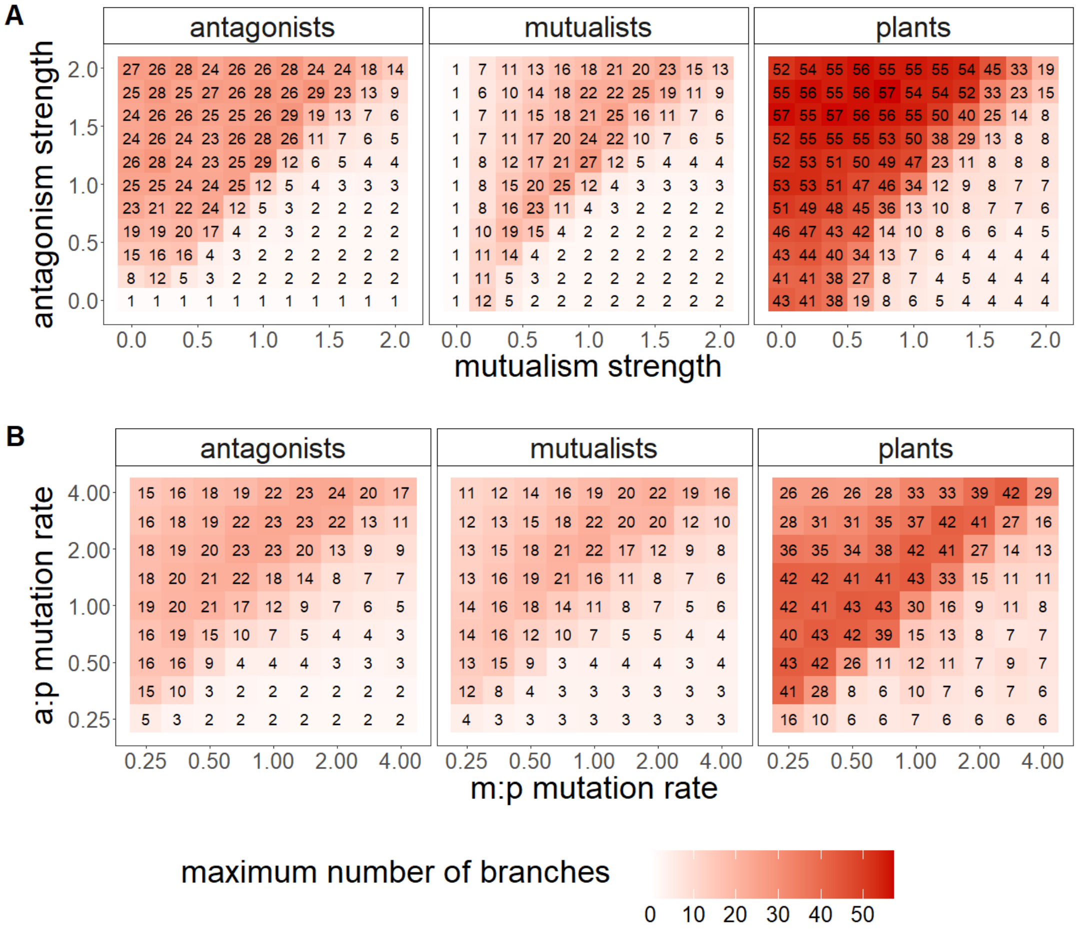

**Figure S3 – Diversification patterns under varying antagonism/mutualism strength and varying speed of evolution.** The plots show the maximum number of branches in each guild for (A) varying maximum strength of mutualism and antagonism ($a_0^{mut}$ and $a_0^{ant}$) and (B) varying relative mutualist and antagonist mutation rates ($\mu_M : \mu_P$ and $\mu_A : \mu_P$; $\mu_P = 2 \cdot 10^{-6}$). A mean is taken over 5 replicates. All other parameters were chosen according to Table 1 from the main text.

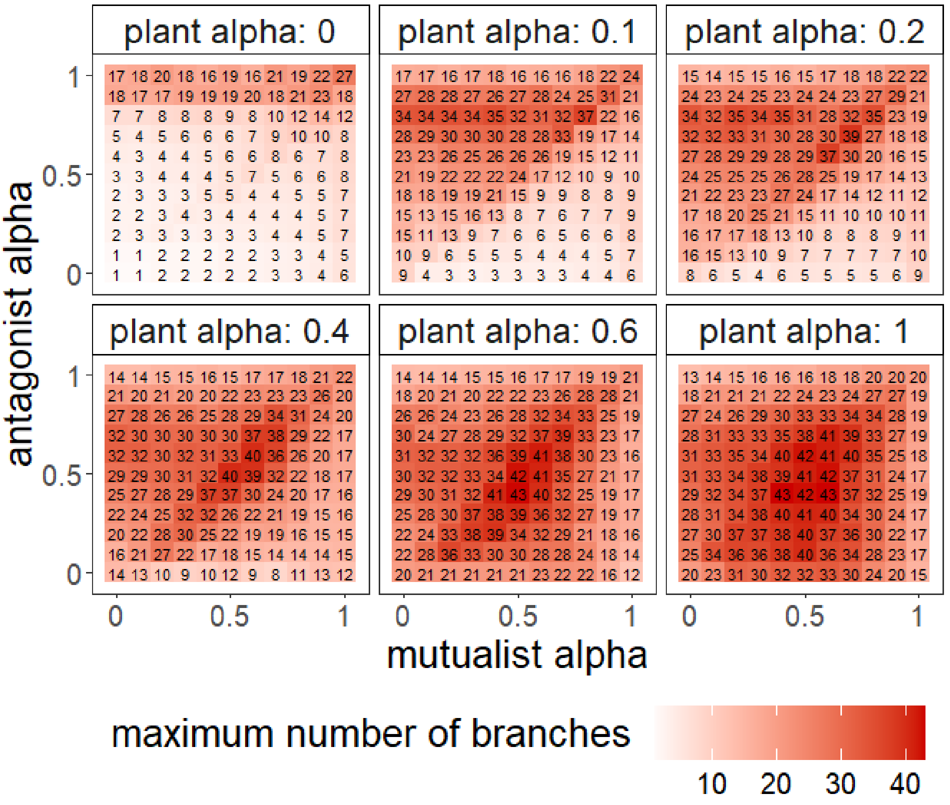

**Figure S4 – Diversification patterns under varying proportion of similarity-based competition.** The plot shows the maximum number of branches for varying shares of similarity-based competition ($\alpha^P$, $\alpha^M$ and $\alpha^A$). A mean is taken over 5 replicates and all guilds. All other parameters were chosen according to Table 1 from the main text.

### S4. Size of extinction avalanches under varying parameter combinations

In this section, we report results on the magnitude of extinction avalanches, complementing the results shown in Figure 1C and D in the main text.

Figure S5 displays the size of extinction avalanches for varying maximum antagonism/mutualism strength and varying relative speed of evolution, broken down by guild. Extinction avalanches in plants are generally larger than those in the other guilds, which follows from the fact that plants diversify more (cf. section S3). Moreover, in all guilds, in the parameter regions where they occur, extinction avalanches tend to be larger if relative strength of antagonism or relative speed of antagonist evolution is high. In these scenarios, avalanche frequency is rather low, in line with the observation from the main text that high avalanche rates are associated with small avalanche size due to less time for prior diversification. The same negative correlation between avalanche size and rate can be observed when varying the shares of similarity-based competition (Fig. S6, compare with Fig. 2 from the main text).

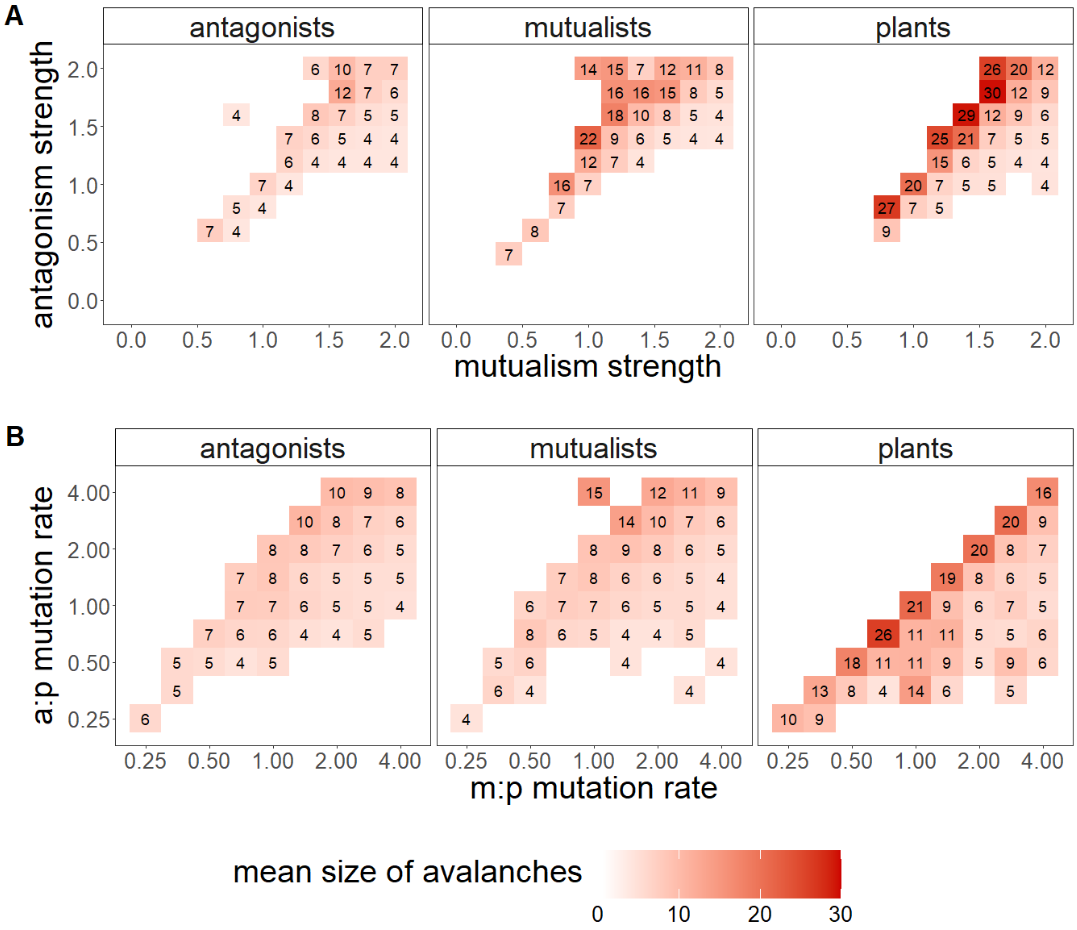

**Figure S5 – Size of extinction avalanches under varying antagonism/mutualism strength and varying speed of evolution, broken down by guild.** The size of extinction avalanches is measured as the maximum number of branches lost within at most 100 million time units. A mean is taken over all avalanches in each guild, using 5 replicates. Avalanche size is computed for (A) varying maximum strength of mutualism and antagonism ($a_0^{mut}$ and $a_0^{ant}$), and (B) varying relative mutualist and antagonist mutation rates ($\mu_M : \mu_P$ and $\mu_A : \mu_P$; $\mu_P = 2 \cdot 10^{-6}$). All other parameters were chosen according to Table 1 from the main text.

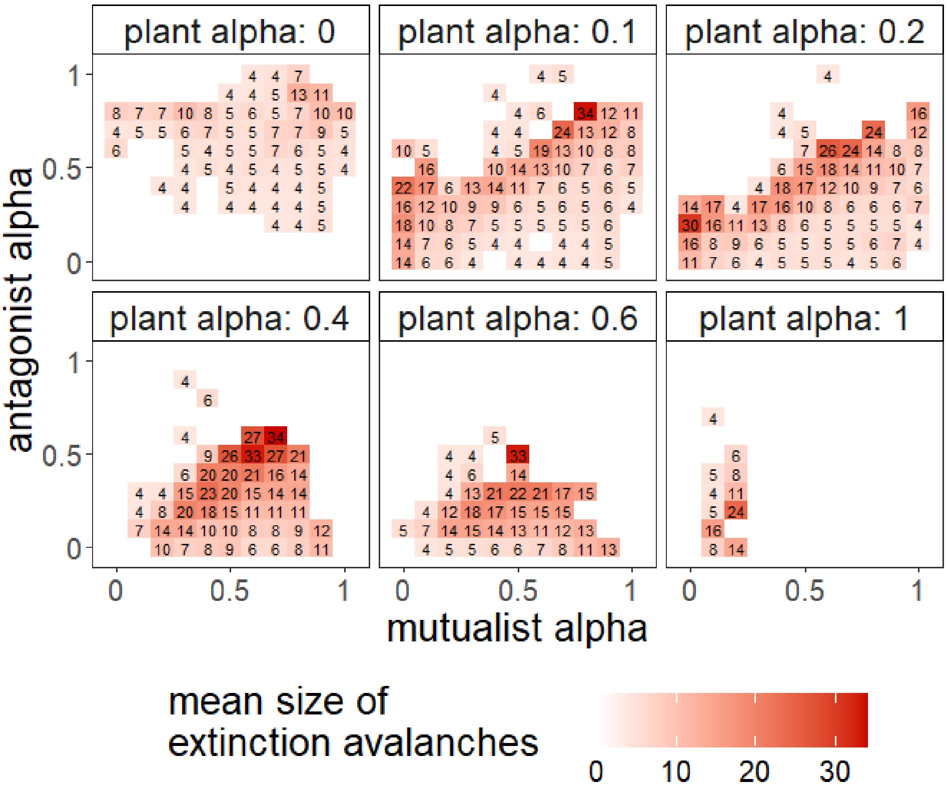

**Figure S6 – Size of extinction avalanches under varying proportion of similarity-based competition**. Plots show the size of extinction avalanches measured as the maximum number of branches lost within at most 100 million time units. A mean is taken over all avalanches in all guilds, using 5 replicates. The mean avalanche size is calculated for varying shares of similarity-based competition ($\alpha^P$, $\alpha^M$ and $\alpha^A$). All other parameters were chosen according to Table 1 from the main text.

### S5. Results from manipulative experiments with multiple avalanches

We repeated the described manipulative experiments (as in Fig. 4 from the main text) with 5 randomly chosen extinction avalanches from simulations with the baseline parametrisation. The results confirm the findings for the example analysed in the main text. Suppressing the evolution of the top or bottom mutualist branch clearly prevents extinctions in four out of five cases (Figure S7A). In the remaining case (avalanche 3 in the figure), an extinction avalanche is highly unlikely even in the baseline scenario, in which all branches are allowed to evolve. Moreover, in all five avalanches, suppressing the evolution of the top or bottom antagonist branch leads to significantly more extinctions than in the baseline scenario. Figure S7B confirms the findings on the ecological mechanisms that cause the extinctions. If the first mutant in an avalanche - always a mutant in an outer mutualist branch - is added to the system at equilibrium, potential subsequent extinctions almost never occur if plant densities are kept constant, but are facilitated if antagonist dynamics are switched off.

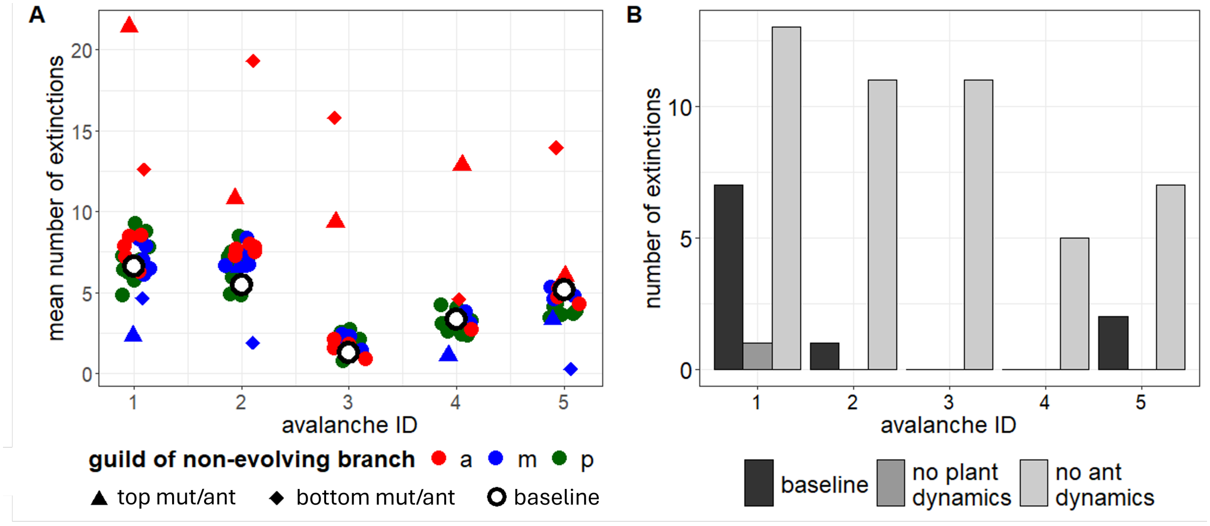

**Figure S7 – Manipulative experiments with 5 example avalanches**. (A) Suppression of evolution of one branch at a time. Simulations are initialised with the state right before the extinction avalanche (calculated as described in section S2). Each point in the figure corresponds to a branch that is prohibited to evolve. The plot shows the total number of extinctions across all guilds within 50 million time units, averaged over 20 replicates. The blue and red triangle (square) correspond to the mutualist and antagonist branch with highest (lowest) trait value. Unfilled circles represent the baseline scenario with all branches evolving. (B) Manipulation of ecological dynamics after mutation, no further evolution. Simulations are initialised with the state right before the first extinction avalanche, but with ecological dynamics at equilibrium and the next (mutualist) mutant added with initial density $N_0$. The plot shows how many branches go extinct after simulating the subsequent ecological dynamics (no evolution) for 100 000 time units in (i) the baseline scenario, (ii) a scenario with constant plant densities, (iii) a scenario with constant antagonist densities.
